# Functionally Mature Bioengineered Human Skeletal Muscle Tissues Capture Essential Aspects of Glucose Metabolism

**DOI:** 10.64898/2026.01.16.698404

**Authors:** Carlos Henriquez-Olguin, Martina Kubec Højfeldt, Christopher Thomas Andrew Lewis, Jesper B. Birk, Tianfang Wang, Zelin Li, Pia Jensen, Pauline Blomquist, Bo Falck Hansen, Roberto Meneses-Valdes, Enrique Toledo, Jeb Hogan, Zhe Wang, Charis-Patricia Segeritz, Jørgen Frank Pind Wojtaszewski, Thomas Elbenhardt Jensen, Anna Blois, Christian Pehmøller, Jonas Roland Knudsen

## Abstract

Human skeletal muscle is a major regulator of whole-body metabolic homeostasis, yet mechanistic insight into human muscle plasticity is limited by the lack of in vitro models with adult-like metabolic and functional maturity. Here, we develop a workflow for generating bioengineered human skeletal muscle tissues that undergo coordinated structural, molecular, and functional maturation and stabilize in an adult-like state by day 21. Time-resolved RNA-seq and protein profiling reveal consolidation of contractile programs alongside progressive metabolic maturation, including increased mitochondrial electron transport chain content, mature mitochondrial network organization, and upregulation of glucose- and glycogen-handling proteins as well as muscle-enriched AMPK isoforms. Functionally, the tissues develop physiological force–frequency behavior, post-tetanic potentiation, and reproducible fatigue responses that are exacerbated by hypoxia and glucose withdrawal. Exercise-like chronic stimulation increases force and endurance with hypertrophy-like remodeling, and these adaptations reverse with detraining. The model also captures pharmacological responsiveness. β2-adrenergic stimulation activates canonical signaling, increases force, limits disuse-related decline, and improves endurance in a glucose-dependent manner. Under physiological insulin and IGF-1 conditions, tissues show robust insulin-stimulated glucose uptake and glycogen synthesis, with punctate glucose transporter 4 (GLUT4) localization. Finally, knockdown of muscle glycogen synthase (GYS1) preserves peak tetanic force but impairs endurance and force recovery under fuel stress, indicating that glycogen metabolism is a key determinant of human muscle resilience.

## 1. INTRODUCTION

Skeletal muscle is central to whole-body homeostasis, accounting for the majority of postprandial glucose uptake [1] and controlling strength, endurance, and metabolic health [2]. Muscle plasticity underlies whole-body adaptations to exercise, physical inactivity, and pharmacological interventions, while muscle maladaptation can lead to insulin resistance, sarcopenia, and multiple chronic diseases [3]. Understanding the molecular pathways that govern human muscle plasticity has clear implications for improving human health, and scientific progress depends on cellular models that faithfully reproduce the physiology of human muscle tissue.

Animal models have been invaluable for dissecting mechanisms of muscle adaptation [4], yet their translational relevance to humans is inherently constrained. Fundamental interspecies differences in fiber-type distribution [5], glycogen storage [6], metabolic regulation [7], and pharmacological responses all limit the predictive value of rodent studies to study human physiology and drug discovery. Human in vivo studies, while critical, face major ethical and technical barriers: permanent damage from experimental interventions must be avoided, genetic perturbations are not feasible, and repeated tissue sampling for mechanistic readouts is challenging [8]. As a result, many central questions in human muscle biology, including determinants of contractile plasticity, pharmacological responsiveness, and the molecular basis of muscle endurance, remain incompletely understood.

Bioengineered human muscle tissues offer a way to bridge this gap. Recent advances have demonstrated the potential of various approaches to bioengineer muscle tissues that recapitulate aspects of myogenic differentiation and contractility [9–20]. In severe myopathies, these platforms have demonstrated the ability to recapitulate disease-associated phenotypes, including abnormal nuclear morphology in differentiated induced pluripotent stem cells (iPSCs) derived from patients with LMNA gene mutations [10], impaired contractile function in Duchenne muscular dystrophy patient-derived differentiated iPSCs compared to differentiated isogenic control cells [21] as well as impaired contractile function in differentiated iPSCs derived from limb girdle muscular dystrophy 2B patients [22]. Certain aspects of adaptations to exercise training, including increased cell size and maximal contractility, have also been replicated by chronic exercise mimicking electrical stimulations [13,19,23,24]. Despite the progress, it remains unclear how accurately these platforms reproduce skeletal muscle metabolism or capture genetic and pharmacological perturbations that modulate muscle bioenergetics. Resolving this is essential because mature human muscle models could provide a uniquely tractable platform for dissecting mechanisms fundamental to metabolic health and drug development.

One area where such models are particularly valuable is glucose metabolism, where mice and humans are profoundly different [25]. Glycogen is a central substrate for endurance performance and metabolic health, and its regulation by glycogen synthase 1 (GYS1) is well established in animal models [26,27]. In humans, patients with Glycogen Storage Disease 0b, characterized by elimination of the *Gys1* gene, are characterized by having low levels of glycogen and poor exercise performance capabilities. However, these patients also present cardiomyopathy already in childhood [28], making interpretation of the role of GYS1 in skeletal muscle based on observations in these patients difficult. Thus, beyond lifelong depletion of GYS1 accompanied by cardiomyopathy and early death, the requirement of GYS1 for skeletal muscle endurance in human models is unknown.

Here, we present a workflow for a human bioengineered muscle tissue model that undergoes profound structural, molecular, and functional maturation and can be subjected to physiological, pharmacological, and genetic interventions that modulate metabolic outputs, including muscle fatigue and glucose uptake. We show that bioengineered human muscle tissues recapitulate native-like plasticity in response to contractile activity and inactivity, β-adrenergic stimulation, and targeted gene silencing. Notably, GYS1 knockdown compromises muscle endurance and recovery, highlighting the limiting role of glycogen metabolism in human muscle contractile function. Together, our findings establish bioengineered muscle tissues as a versatile model to dissect mechanistic regulators of human muscle biology, bridging the translational gap between animal studies and clinical research.

## 2. RESULTS

### 2.1 Bioengineered human muscle tissues reach a stable structural and molecular mature state

To generate metabolically and functionally mature human muscle, primary myoblasts were embedded in a fibrinogen-based hydrogel, cast into 24-well plates, and differentiated for up to 35 days (Fig. 1A), allowing morphological, transcriptomic, and functional characterization across maturation. By day 21, confocal imaging revealed aligned, multinucleated fibers exhibiting peripheral nuclear orientation and organized sarcomeres (Fig. 1B-C), closely resembling the architecture of adult skeletal muscle [31,32]. Comparison of bioengineered muscle tissues at day 21 with conventional 7-day monolayer myotubes revealed a strong induction of sarcomeric and contractile genes, including MYBPC1, MYL1, MYH2, and other canonical muscle markers (Fig. 1D). Pathway enrichment analysis of the upregulated gene set showed significant enrichment for muscle contraction and muscle system processes, while downregulated genes were associated with RNA processing and translation (Fig. 1E).

**Figure 1.**
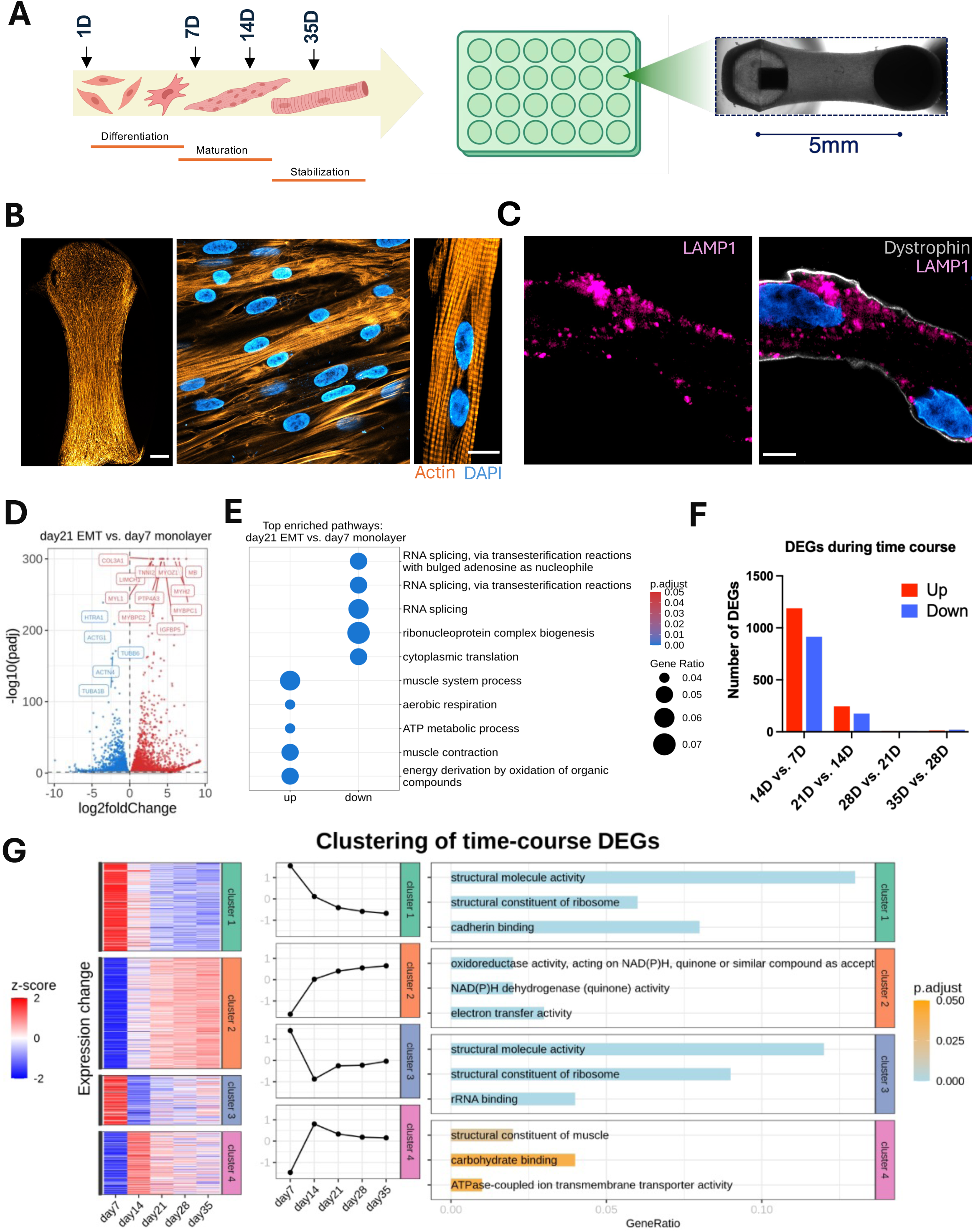
Bioengineered human muscle tissues undergo structural and molecular maturation. **(A)** Schematic of the muscle tissue bioengineering workflow, in which primary human myoblasts are embedded in fibrin-based hydrogels, cast in 24-well plates, and differentiated for up to 35 days. **(B)** Confocal microscopy images of day-21 bioengineered muscle bundles showing aligned myofibers, multinucleated structures (DAPI, blue), and sarcomeric actin (orange hot). **(C)** High-resolution confocal images of day-21 bundles showing LAMP1 (lysosomes) and Dystrophin (plasma membrane) staining. **(D)** Volcano plot comparing differentiated bioengineered muscle tissues with 7-day 2D monolayer myotubes **(E)** Gene ontology enrichment of upregulated DEGs showing categories associated with muscle system processes. **(F)** Number of differentially expressed genes (DEGs) across the time course from day 7 to day 35, with the largest shift observed between maturation time-course. **(G)** Clustering of time-course DEGs identified distinct maturation-associated transcriptional programs. (n=3-4 for each group). EMT = Bioengineered human muscle tissues, Monolayer = myotubes cultured in monolayer, DEG = differentially expressed genes. Scale bars: B, 500 µm (right) and 10 µm (left); C, 10 µm.

Temporal RNA-seq from day 7 to day 35 showed extensive transcriptomic remodeling early in differentiation, with more than 2,000 genes differentially expressed between days 7 and 14 (Fig. 1F), marking the peak of transcriptional change. By day 21, this number dropped to 420 compared to day 14, and from day 21 onward, the bioengineered human muscle tissue remained transcriptionally stable, with only minor changes detected (Fig. 1F). Clustering of time-course DEGs revealed distinct maturation trajectories (Fig. 1G). Genes that increased gradually over time (cluster 2), such as SDHA, were enriched for mitochondrial oxidoreductase and electron transfer functions, indicating progressive metabolic maturation. In contrast, genes that rose by day 14 and remained relatively stable thereafter (cluster 4), including DMD, were enriched for structural components of muscle, reflecting consolidation of contractile architecture. Additional clusters captured early differentiation signatures (cluster 1) and genes downregulated with maturation (cluster 3), together outlining a coordinated shift from differentiation programs toward stable structural and metabolic specialization.

Overall, bioengineered muscle tissues reach a stable, adult-like state by day 21, defined by mature structural and metabolic programs. This maturation-mediated transcriptional stability provides a robust platform for longitudinal mechanistic studies of human muscle biology.

### 2.2 Maturation of bioengineered human muscle tissues exhibits coordinated structural, contractile, and metabolic maturation

To validate muscle maturation at the protein level, we used our RNA-seq dataset to nominate maturation markers and generated an independent set of tissues for western blot analysis across the time course. We first examined canonical myogenic regulators. RNA-seq showed that these factors peaked early during differentiation: MYF5 became undetectable after day 7, MYOD1 and MYOG decreased after day 7, and MYF6 increased (Fig. S1A). Western blotting mirrored these trends, with MYOD1 detectable at day 2 but absent at later time points, and myogenin showing a ∼50% decrease from day 2 to day 23 that did not reach statistical significance (Fig. S1B).

We next profiled muscle gene markers based on [33] categorized into fast-twitch genes, slow-twitch genes, fiber-type–independent structural genes and genes not expressed in mature skeletal muscle. We compared expression in 7-day monolayer myotubes with bioengineered tissues differentiated for 7 or 21 days. Overall, slow-twitch genes were higher in tissues than in monolayer cultures, with some markers peaking at day 7 and others increasing further by day 21 (Fig. 2A). All fast-twitch markers were upregulated in tissues at both time points vs. monolayer myotubes, with most of them higher in expression at day 21 than at day 7 (Fig. 2A). The fiber-type-independent markers also indicated highest muscle maturity after 21 days in bioengineered tissues (Fig. 2A), while the muscle markers not expressed in mature skeletal muscle were lowest at day 21 (Fig. 2A). Overall, these data clearly indicate improved skeletal muscle maturity in the bioengineered muscle tissue with highest maturity at the 21-day timepoint and revealed that particularly fast-twitch fiber maturation seems to be enhanced in the tissues vs. monolayer culture.

**Figure 2.**
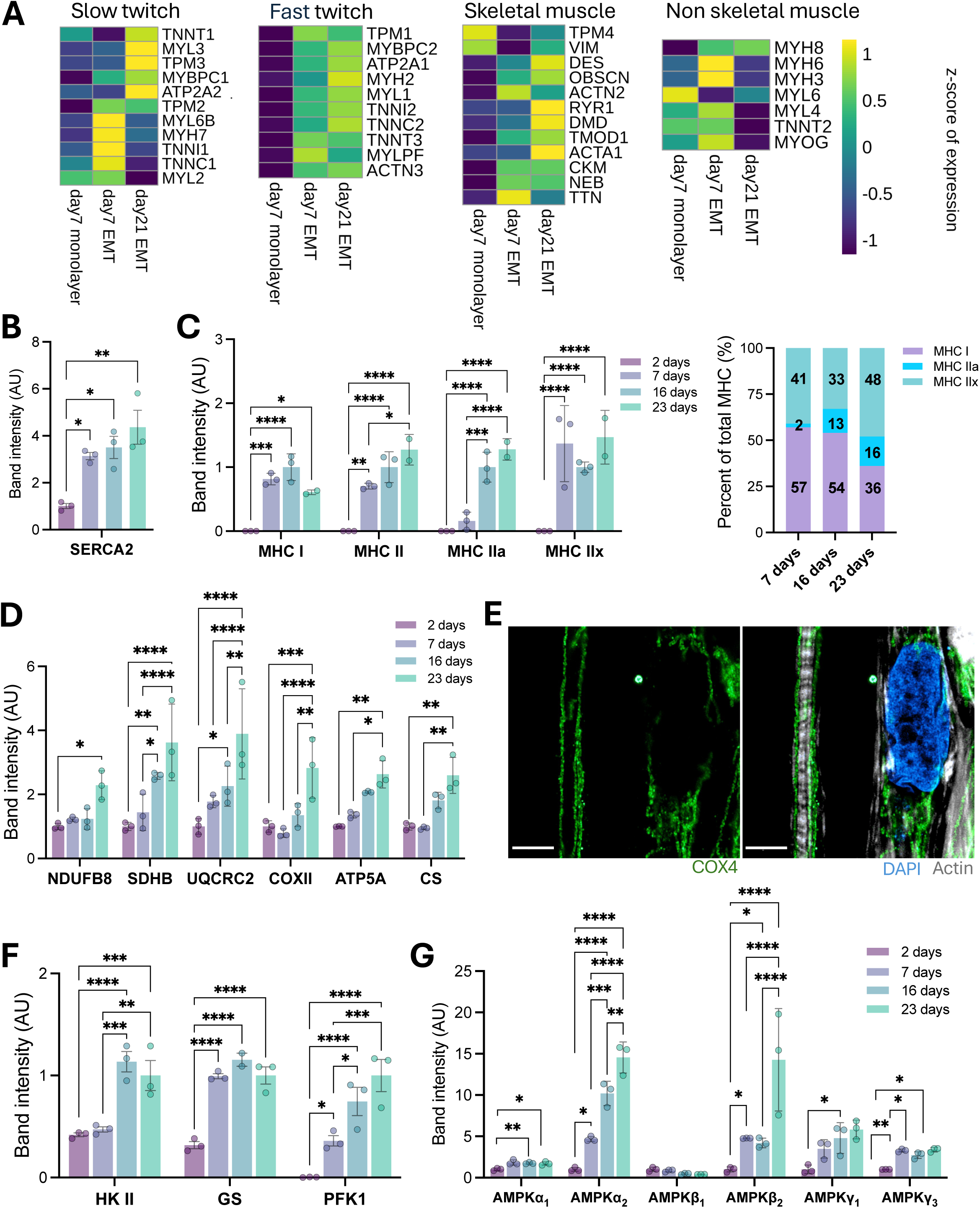
Muscle-related gene and protein expression in bioengineered human muscle tissues. **(A)** Heatmaps showing expression of slow-twitch, fast-twitch, general skeletal muscle, and non-skeletal muscle gene sets in bioengineered tissues at day 7 and day 21 compared with 7-day 2D monolayer myotubes. **(B)** SERCA2 protein levels measured across differentiation time points. **(C)** Protein abundance and relative proportion of myosin heavy chain isoforms. **(D)** Protein abundance of mitochondrial electron transport chain subunits from complexes I–V, including NDUFB8, SDHB, UQCRC1, COX4, ATP5A, and citrate synthase (CS). **(E)** AiryScan confocal images of COX4 staining (green) and actin (grey) in muscle tissues. **(F)** Quantification of metabolic enzymes including hexokinase II, glycogen phosphorylase, and phosphorylase kinase at indicated time points. **(G)** Protein expression of glycolytic enzymes and AMPK subunits, including AMPKα2, AMPKβ2, AMPKγ1, and AMPKγ3. Corresponding gene-expression data and representative western blots are shown in Fig. S2C–D. Data (n=3) are presented as mean ± SEM, with individual values shown where applicable. Statistical comparisons were performed using one-way or two-way ANOVA with appropriate multiple-comparisons correction. \**p* < 0.05, \*\**p* < 0.01, \*\*\**p* < 0.001, \*\*\*\**p* < 0.0001. EMT = Bioengineered human muscle tissues, Monolayer = myotubes cultured in monolayer. Scale Bar =5 mm.

The pattern in fiber-type changes was also confirmed at the protein level. The slow-twitch markers SERCA2 and MHCI increased drastically between day 2 and 7 before reaching more stable expression levels (Fig. 2B-C). The fast-twitch marker MHCII continued to increase beyond day 7, driven by an early increase in MHCIIx and a later increase in MHCIIa (Fig. 2C). After 23 days of differentiation, 36% of the myosin was composed of slow twitch while 64% represented fast twitch (16% IIa and 48% IIx) (Fig. 2C). Further protein analyses supported the progressive metabolic maturation observed in Fig. 1, confirming increased abundance of mitochondrial electron transport chain subunits (Fig. 2D) and maturation of mitochondrial network architecture, including prominent subsarcolemmal and perinuclear localization (Fig. 2E). Proteins involved in glucose uptake and glycogen metabolism also increased with maturation (Fig. 2F). Furthermore, the expression of the muscle dominant α2, β2, γ1 and γ3 isoforms of the heterotrimeric energy sensor AMPK were upregulated (Fig. 2G). Gene data for the proteins shown in Fig. 2D-G, as well as representative western blots are shown in Fig. S1C-D.

Overall, this profiling of the bioengineered human muscle tissues clearly revealed an initial immature differentiation that then appeared diminished within the 7-14 days that cells are allowed to mature in most in vitro muscle platforms. However, interestingly beyond the 14 days of differentiation, the structural and contractile maturation continued and was complemented by molecular and structural markers of metabolic maturation. By day 21, bioengineered human muscle tissues exhibited a plethora of hallmarks of skeletal muscle maturation, including aligned multinucleated fibers with striated sarcomeric organization, a stable transcriptional profile enriched in contractile and metabolic markers at both the gene and protein levels.

### 2.3 Physiologically relevant force and fatigue profiles in bioengineered muscle tissues

We next assessed whether these maturation-associated molecular changes translated into functional output by quantifying tissue contractility. Electrically evoked force increased in parallel with the transcriptional maturation of the tissues. Twitch force rose from ∼280 μN at day 7 to ∼1100 μN by day 21 for twitch and from ∼750 μN at day 7 to ∼1,800–2,000 μN after day 14 for tetanic contractions (Fig. 3A). The increased force was accompanied by maturation of contractile kinetics as contraction and relaxation speed progressively increased while the contraction and relaxation times were unchanged (Fig. S2A). From around day 14 and onwards the maximal force production was stable for the duration of the experiments (Fig. 3B) and tissues exhibited physiologically relevant frequency-dependent force development, transitioning from twitch-like to tetanic contractions at 1-10 Hz (Fig. 3C). The 3D muscles also responded to fiber-type specific modulation of the contractile machinery. Slow-twitch myosin inhibitor Mavacamten [34,35] and pan-myosin inhibitor Blebbistain [36,37] both decreased contractility while fast-twitch troponin activator Tirasemtiv [38,39] evoked increased force production (Fig. S2B). Having established contractile maturity, we next studied whether post-tetanic potentiation, a phenomenon of increased twitch force production observed following a prior tetanic contraction either evoked electrically or voluntary [40–42], was observable in the bioengineered muscle tissue. Indeed, prior tetanic stimuli (Fig. S2C) increased twitch and summed twitch force in the tissues (Fig. 3D & S2D).

**Figure 3.**
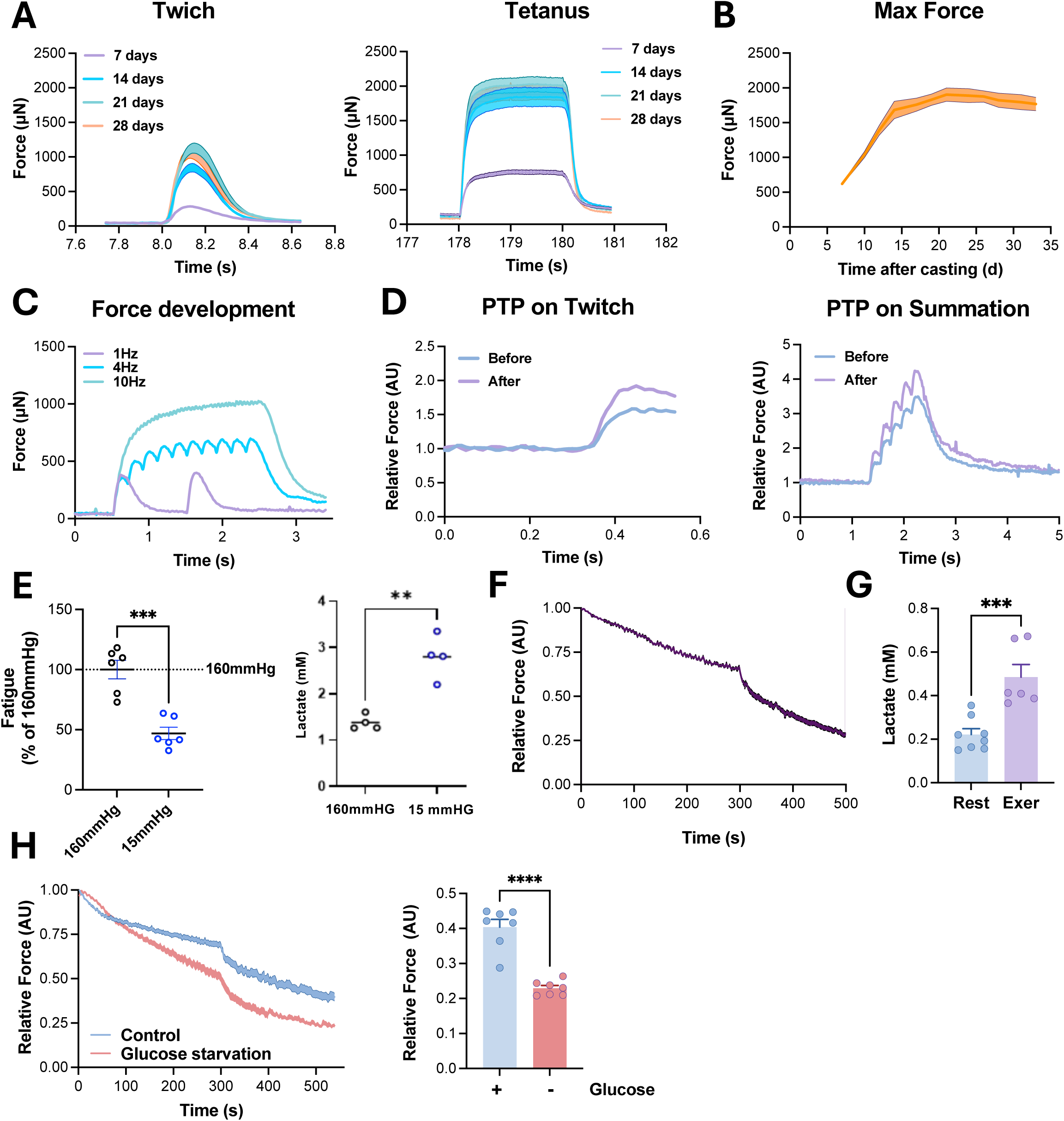
Contractile properties, fatigue behavior, and metabolic responses of bioengineered human muscle tissues. **(A)** Electrically evoked twitch and tetanic force traces from tissues differentiated for 7, 14, 21, and 28 days. **(B)** Maximal tetanic force recorded from day 7 to day 28 after casting **(C)** Force–frequency relationship showing twitch responses and transition to tetanic contractions at stimulation frequencies between 1 and 10 Hz. **(D)** Post-tetanic potentiation (PTP) assessment. Left: representative twitch force traces before and after a tetanic stimulus. Right: averaged relative twitch force before and after potentiation. **(E)** Fatigue protocol performed under normoxic (∼160 mmHg O₂) and hypoxic (∼15 mmHg O₂) conditions. Left: fatigue development expressed as percentage of force remaining. Right: lactate concentrations measured after contractions under each condition (n=4-6). **(F)** Time course of force decline during an optimized fatigue protocol (protocol 1) **(G)** Lactate concentrations measured in resting versus contracting tissues following the fatigue protocol (n=6-8). **(H)** Fatigue development under glucose and glucose-free conditions during protocol 1 stimulations. Left: relative force during repeated contractions. Right: percent force remaining after the protocol (n=7). Data are presented as mean ± SEM, with individual values shown where applicable. Statistical comparisons were performed using one-way or two-way ANOVA with appropriate multiple-comparisons correction. \*\**p* < 0.01, \*\*\**p* < 0.001, \*\*\*\**p* < 0.0001.

To characterize the fatiguability of the bioengineered human muscle tissue and its dependence on oxidative phosphorylation, we performed an extensive fatiguing contraction protocol (Fig. S2E) during normoxia (∼160mmHg O_2_) or hypoxia (∼15mmHg O_2_). As expected, hypoxia accelerated fatigue development and increased lactate production (Fig. 3E). Subsequently, we optimized the contraction protocol to better mimic the protocols used in excised muscle from rodents or humans [40,43,44]. This contraction regimen consistently caused a gradual fatigue development in the tissues down to ∼30-40% of the starting force after 8-9 min (Fig. 3F) and caused lactate accumulation compared to resting tissues (Fig. 3G). Removing glucose from the cell culture medium further accelerated fatigue development (Fig. 3H).

Overall, these findings demonstrate that bioengineered human muscle tissues exhibit contractile features mimicking those reported from in situ/ex vivo studies in adult muscle fibers in rodents and humans. Furthermore, the tissues respond to physiological modulators such as prior contractions, reduced oxygen and glucose availability.

### 2.4 Contraction training drives structural remodeling and metabolic reprogramming

To assess whether bioengineered muscle tissues recapitulated exercise training responses, we subjected them to repeated contractile stimulation bouts. Tetanic stimulation (50Hz, 100mAmp, 5 sec every 50 sec) took place 3 times weekly, starting with 30 min stimulation the first day, increasing to 60 min the second day and then finally 90 min stimulation which was then continued throughout the experiment. As predicted, stimulation increased maximal force production by ∼14% (Fig. 4A) and was accompanied by marked remodeling as confocal imaging revealed a visibly larger myofibrillar density and myotube size in trained tissues (Fig. 4B), consistent with structural adaptations observed during in vivo muscle hypertrophy. Importantly, these morphological changes were accompanied by expected molecular remodeling, as protein expression of GLUT4 and PDH-E1α, proteins well established to be regulated by exercise training in humans [45,46], increased by ∼59 and ∼43% respectively (Fig. 4C).

**Figure 4.**
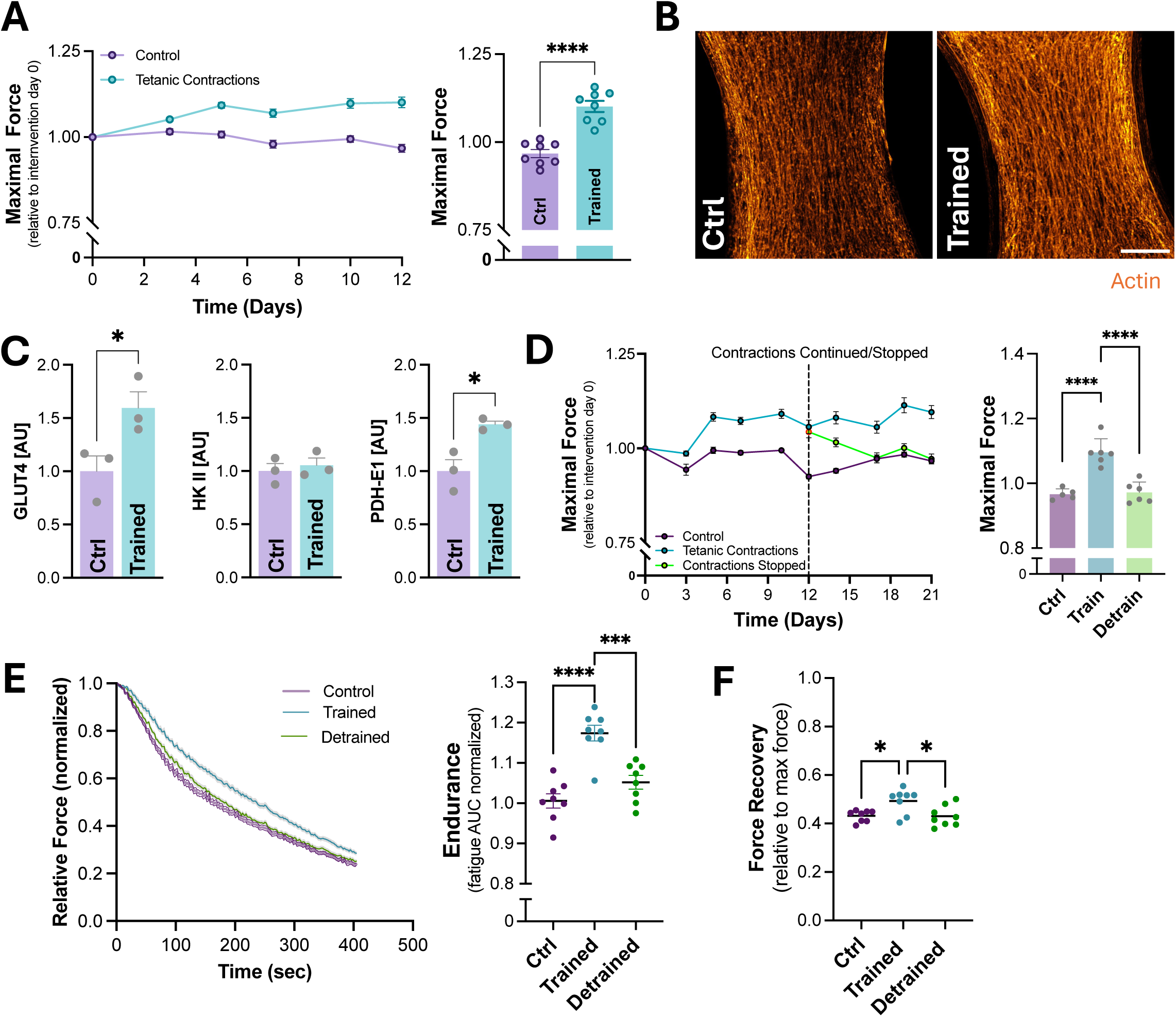
Exercise-like contractile training drives structural remodeling and improves force and endurance in bioengineered human muscle tissues, with reversibility upon detraining. **(A)** Time course of maximal tetanic force during a 12-day electrical training regimen (left) and endpoint quantification of maximal force in control versus trained tissues (right). **(B)** Representative confocal images showing actin organization in control and trained tissues **(C)** Protein abundance of exercise-responsive metabolic markers in control versus trained tissues, including GLUT4, HKII, and PDH-E1α. **(D)** Maximal force overtime in control tissues, continuously trained tissues, and tissues in which training was stopped after the training period (left; dashed line indicates the time point when contractions were continued or withdrawn), with endpoint comparison of maximal force between groups (right). **(E)** Fatigue profiles following contractions (protocol 2) (left) and quantification of endurance as force decline in control, trained, and detrained tissues (right). **(F)** Recovery of force after the fatigue protocol expressed relative to maximal force in control, trained, and detrained tissues. Data are presented as mean ± SEM, with individual values shown where applicable. Statistical comparisons were performed using t-student, one-way or two-way ANOVA with appropriate multiple-comparisons correction. \**p* < 0.05, \*\**p* < 0.01, \*\*\**p* < 0.001, \*\*\*\**p* < 0.0001.

To evaluate the persistence and reversibility of training-induced changes, we performed a detraining experiment in which tissues were first matured for 3 weeks and then trained for 12 days, followed by withdrawal of stimulation in one group. During the training phase, maximal tetanic force increased by ∼14% in stimulated tissues while withdrawal of stimulation led to a progressive decline in force output, with values returning to untrained levels by day 21 (Fig. 4D), indicating that the training-induced gains were reversible.

Assessment of the fatigue profile following the 21 days with or without training and detraining also revealed reversible improvements in the endurance during the contraction reflected by larger force decline during the stimulation (Fig. 4E) as well as an improved force recovery after contraction cessations in the trained vs. non-trained and detrained tissues (Fig. 4F). Together, these results demonstrated that bioengineered human muscle tissues recapitulate key structural and metabolic features of *in vivo* muscle training and exhibited reversible adaptations. Training-induced improvements in force generation and endurance were lost upon detraining, underscoring the dynamic molecular plasticity of these tissues.

### 2.5 Bioengineered human muscle tissues respond to β_2_-adrenergic stimulations

To assess pharmacological responsiveness, we next focused on β2-adrenergic receptor (β2AR) signaling, a well-established regulator of human muscle plasticity. We therefore used the β2AR agonist clenbuterol as a pharmacological probe. Acute treatment (20 min) with 10 nM clenbuterol activated canonical downstream phosphorylation events, including phosphor(p)-HSL (Ser660), p-CREB (Ser133), and p-RPS6 (Ser235/236) (Fig. 5A), while chronic treatment elevated B2AR responsive genes (*PPARGC1A, NR4A3, NR4A1, PGK1, GYS1*) (Fig. 5B). Functionally, clenbuterol induced a dose-dependent increase in maximal tetanic force, with a ∼25% enhancement observed at 10 nM (Fig. 5C). Morphological analysis revealed myofibrillar thickening and enhanced sarcomere alignment, recapitulating the adaptations observed under tetanic training (Fig. 5D).

**Figure 5.**
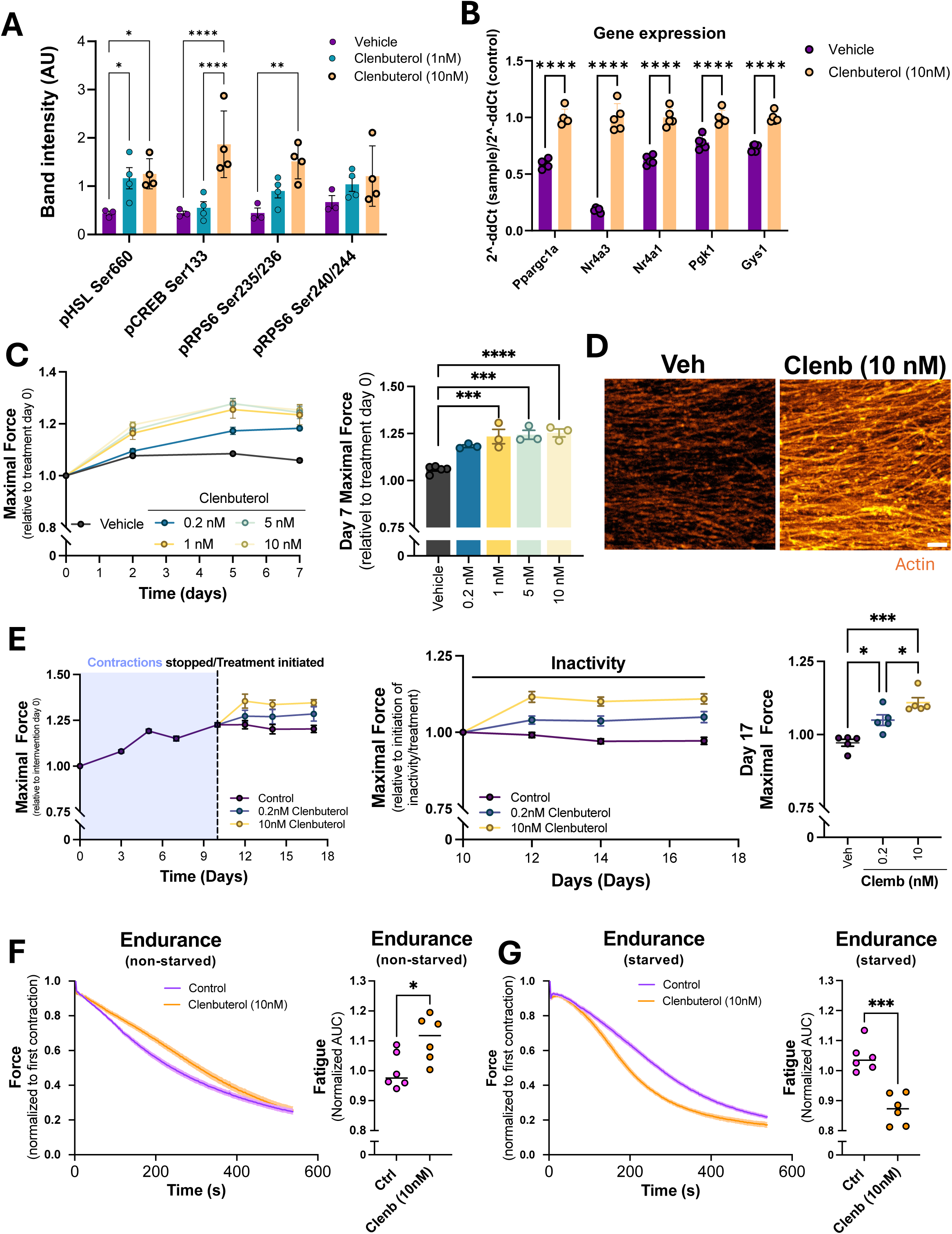
β_2_-adrenergic signaling effects on bioengineered human muscle tissues. **(A)** Effect of acute treatment with clenbuterol (10 nM, 20 min) on phosphorylation of β-adrenergic (B2AR) downstream targets including HSL (Ser660), CREB (Ser133), and RPS6 (Ser235/236) (n=4). **(B)** Gene expression of B2AR–responsive transcripts (*PPARGC1A*, *NR4A3*, *NR4A1*, *PGK1*, *GYS1*, and others) after chronic clenbuterol treatment at the indicated doses (n=5). **(C)** Maximal tetanic force measured over time in tissues exposed to increasing clenbuterol concentrations (0.2–10 nM). Right: day-7 maximal force relative to vehicle for each dose (n=3). **(D)** Representative confocal images of myofibrillar structure in vehicle- and clenbuterol-treated tissues (n=2) **(E)** Maximal tetanic force during an inactivity protocol with or without clenbuterol treatment. Left: force traces over the detraining period. Right: day-7 maximal force in vehicle versus clenbuterol-treated groups (n=5). **(F)** Fatigue development during repeated contractions (protocol 2) in glucose-containing media in control versus clenbuterol-treated tissues. Inset: endurance values at the end of the protocol (n=5). **(G)** Fatigue development during repeated contractions (protocol 2) in glucose-free media in control versus clenbuterol-treated tissues (n=5). Data are presented as mean ± SEM, with individual values shown where applicable. Statistical comparisons were performed using one-way or two-way ANOVA with appropriate multiple-comparisons correction. \*\**p* < 0.01, \*\*\**p* < 0.001, \*\*\*\**p* < 0.0001.

Importantly, we tested whether clenbuterol could protect against inactivity-induced functional decline. Intriguingly, 7 days of clenbuterol treatment improved force production in detrained tissues by 8% at submaximal (0.2nM) and 14% at maximal (10nM) doses (Fig. 5E). Finally, due to the positive effect of B2AR agonists on anaerobic muscle performance [47,48] and the reprogramming towards reliance on glucose utilization in the muscles seen following clenbuterol treatment [49], we assessed fatigue development in the tissues with and without glucose supplemented in the cell culture medium. Intriguingly, similar to what is reported in humans, B2AR agonism by clenbuterol improved endurance in the presence of glucose (Fig. 5F) while glucose starvation impaired endurance (Fig. 5G). These findings identified B2AR signaling as a potent modulator of bioengineered muscle plasticity and a pharmacological strategy to counteract disuse-induced atrophy. Furthermore, the dose-responsiveness and the differential effect of the drug seen with and without glucose in the medium align with observations in humans *in vivo*, highlighting the strong pharmacological translatability of this platform.

### 2.6 Bioengineered human muscle tissues exhibit GLUT4 expression and mature insulin responsiveness

To study insulin responsiveness, we screened medium formulations with reduced insulin and IGF-1 concentrations mimicking physiological levels. In 7-day differentiated myotubes, the selected medium formulation enhanced insulin sensitivity at the level of pAkt compared to standard medium (EC_50_ =0.67 vs. 11.52nM) (Fig. S3A). Furthermore, it shifted glucose transporter expression, reducing GLUT1 and GLUT3 while increasing GLUT4 expression (Fig. S3B), consistent with mitigation of the culture-induced insulin resistance typically seen in primary human muscle cells [50]. Intriguingly, the metabolically-favorable mediums robustly increased GLUT4 protein expression in bioengineered muscle tissues (Fig. 6A). Immunofluorescence and high-resolution imaging revealed punctate GLUT4 localization throughout the tubes (Fig. 6B), consistent with the intracellular distribution reported in mature skeletal muscle [51–54].

**Figure 6.**
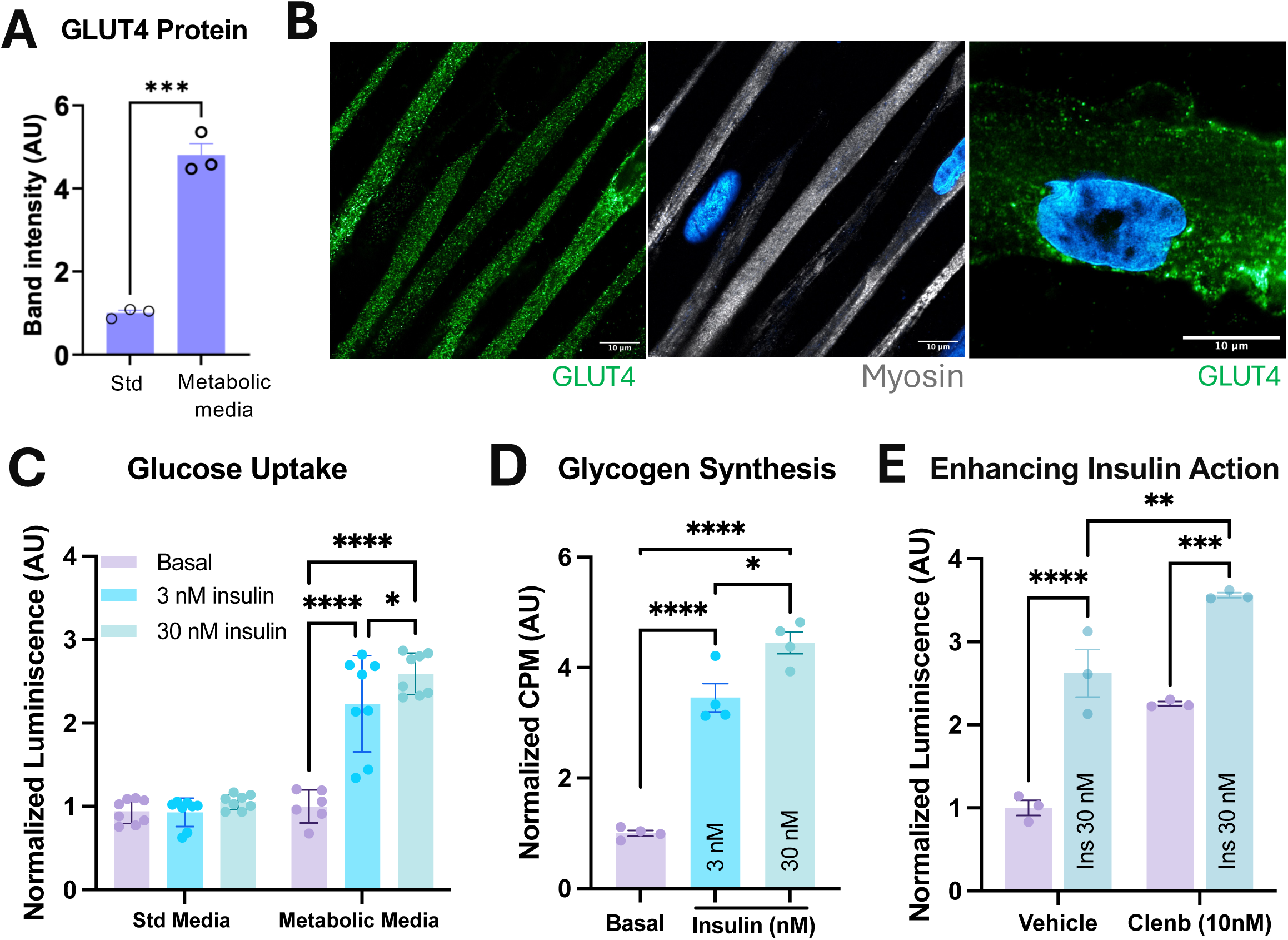
GLUT4 expression and insulin responsiveness in bioengineered human muscle tissues. **(A)** GLUT4 protein abundance measured in tissues maintained in standard versus physiologically modified metabolic media (n=3). **(B)** Representative confocal images showing GLUT4 localization in bioengineered muscle tissues. Left: GLUT4 (green) and myosin (orange) distribution along the fiber. Right: higher-magnification image showing punctate GLUT4 signal in the perinuclear compartment (n=3). **(C)** Glucose uptake under basal conditions and after stimulation with 3 nM or 30 nM insulin in standard or metabolic media (n=8). **(D)** Glycogen incorporation measured under basal conditions and following stimulation with 10 nM or 30 nM insulin (n=4). **(E)** Glucose uptake in basal and insulin-stimulated conditions following treatment with vehicle or 10 nM clenbuterol and 30 nM insulin (n=3). Data are presented as mean ± SEM, with individual values shown where applicable. Statistical comparisons were performed using one-way or two-way ANOVA with appropriate multiple-comparisons correction. \**p* < 0.05, \*\**p* < 0.01, \*\*\**p* < 0.001, \*\*\*\**p* < 0.0001.

Upon insulin stimulation, glucose uptake increased by ∼125% at 3nM and ∼160% at 30nM insulin in the optimized medium (Fig. 6C), demonstrating strong responsiveness and sensitivity comparable to mature muscle ex vivo [55–57]. Consistently, glycogen synthesis increased in a dose-dependent manner in response to insulin (Fig. 6D). Finally, stimulation with 10 nM clenbuterol further increased both basal and insulin-stimulated glucose uptake (Fig. 6E), highlighting the translatability to human in vivo physiology also on this readout [58,59]. Altogether, these data demonstrate that lowering insulin and IGF-1 to physiological levels further improved the metabolic maturity of the tissues and resulted in insulin-responsive tissues.

### 2.7 GYS1 is required for endurance and metabolic adaptation

Finally, to examine the role of GYS1 in regulating muscle endurance, we RNA silenced GYS1 in bioengineered human muscle tissues. Knockdown effectively reduced both GYS1 gene (∼90%) and protein (∼60%) levels (Fig. 7A-B), confirming robust suppression. Representative western blots are shown in Fig. S3C. Interestingly, GYS1 deficiency did not impair maximal force production throughout the experiment (Fig. 7C). Muscle tissue endurance under standard glucose conditions was also unaffected (Fig. 7D), albeit a small but significant impairment was observed during recovery following the contraction bout (Fig. 7E). However, under glucose-starved conditions designed to increase glycogen reliance, GYS1-deficient tissues showed significantly impaired muscle endurance (Fig. 7F) as well as impaired force production in the recovery following the contraction bout (Fig. 7G). These findings indicate that GYS1 is dispensable for peak tetanic force production but is required to sustain contractile function under energetic/fuel stress, underscoring a key role for glycogen metabolism in metabolic resilience.

**Figure 7.**
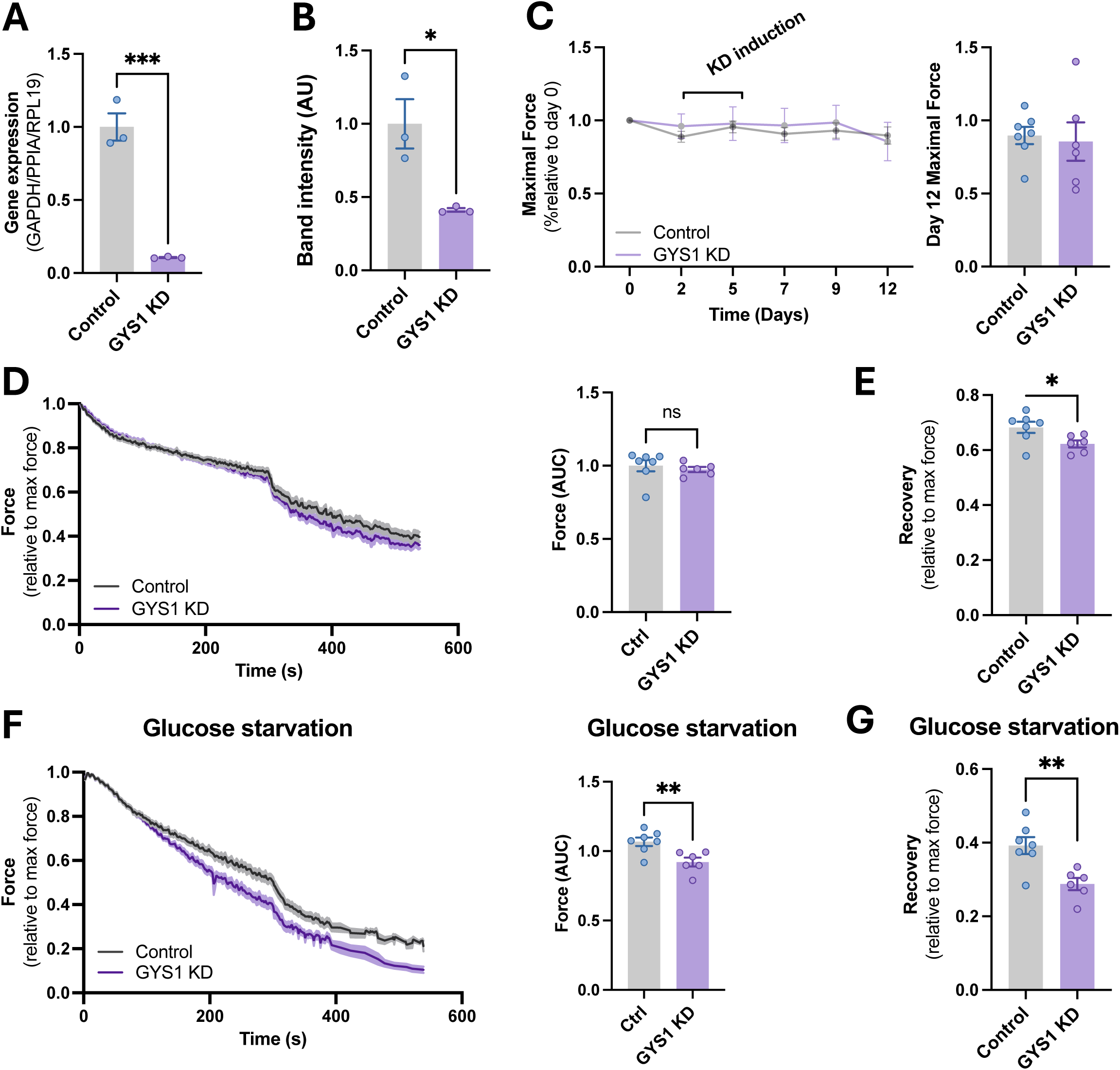
Identification of GYS1 as important for endurance in bioengineered human muscle tissues. **(A)** GYS1 mRNA levels in control and GYS1 knockdown tissues (n=3). **(B)** GYS1 protein abundance in control and GYS1 knockdown tissues (n=3). **(C)** Maximal tetanic force measured over the course of the experiment after knockdown induction (left) and maximal force on day 12 (right) (n=6). **(D)** Fatigue development during repeated contractions (protocol 1) under standard glucose conditions (left) and relative force at the end of the protocol (right) (n=6-7). **(E)** Recovery force following the fatigue protocol under standard glucose conditions, shown relative to maximal force (n=6-7). **(F)** Fatigue development (protocol 1) under glucose-starved conditions (left), relative force at the end of the protocol (middle), and force measured during glucose starvation on day 10 (right) (n=6-7). **(G)** Recovery force following the glucose-starvation fatigue protocol, shown relative to maximal force (n=6-7). Data are presented as mean ± SEM, with individual values shown where applicable. Statistical comparisons were performed using t-student, one-way or two-way ANOVA with appropriate multiple-comparisons correction. \**p* < 0.05, \*\**p* < 0.01, \*\*\**p* < 0.001, \*\*\*\**p* < 0.0001.

## 3. DISCUSSION

Our study provides evidence that bioengineered human skeletal muscle tissues can recapitulate key structural, molecular, and functional features of native muscle and respond predictably to physiological, pharmacological, and genetic perturbations. Furthermore, we identified a conditional requirement in human muscle tissue endurance, consistent with its presumed importance to human contractile performance in vivo. This work thus offers both methodological advancements for translational muscle modeling and new biological insights into the role of human glycogen metabolism.

One key observation in the current study is that bioengineered muscle tissues exhibited progressive and coordinated maturation over three weeks, reflected in structural organization, contractile output, and transcriptional stability. Confocal imaging revealed aligned, sarcomeric fibers and mitochondrial organization, while RNA-seq demonstrated upregulation of gene programs involved in contraction, oxidative metabolism, and myogenesis. By day 21, both force output and gene expression had stabilized, indicating a plateau of maturation and supporting long-term experimental applications starting from day 21. While the tissue maturation plateaued after 3 weeks, importantly, profound maturations of both structural, contractile and metabolic processes were seen beyond 7-14 days of differentiation. At the protein level, notably both contractile fast-twitch and metabolic proteins continued to increase beyond this time point. Furthermore, some muscle development markers such as embryonic myosin and Myogenin were increased at 7 days in the tissues vs. 7 days monolayer culture, but then markedly dropped at later time points below both the levels in tissue and monolayer cultures at day 7. Altogether, these data emphasize the potential of this model to study molecular processes relevant for mature muscle biology with genetic and pharmacological interventions/perturbations after the initial confounding period with high myogenic activity.

Another key observation was that the bioengineered muscle tissues showed robust and translatable responses to external stimuli. Both acute interventions, such as hypoxia, glucose starvation, myosin and troponin modulations and insulin stimulation, as well as chronic interventions including training, detraining and β2AR stimulation were all recapitulated well in the model. To our knowledge, this is the first time acute physiological interventions such as hypoxia and glucose starvation is reported to impact fatigue development in a human in vitro model and also the first time that exercise training-mimicking stimulations is demonstrated to be reversible, adding important translatability validation of bioengineered muscle tissues, likely to be preserved across the growing number of these models [9,10,12,14,15,17–20,23,60].

Repeated contractile stimulation mimicking exercise training induced marked improvements across all measured parameters: increased force generation, enhanced endurance, and upregulation of endurance-exercise responsive metabolic proteins such as GLUT4 and PDH-E1α. These adaptations were reversible upon cessation of stimulation, mirroring the detraining responses observed *in vivo*. These observations build on elegant work by Khodabukus et al., [19] that reported both increased cross-sectional area, structural improvements as well as increased force production following 1 week of electrical pulse stimulation after one week of differentiation of tissue-engineered myobundles. However, in contrast to us, Khodabukus et al. did not observe any changes in GLUT4 expression with either 1Hz or 10Hz stimulations. This may be due to different stimulation timing (after 1 week vs. after 21 days) or the stimulation protocol itself (continuous stimulation for 1 hour every 7 hours for 1 week vs. 90min per day 3 days a week). As increased GLUT4 expression is a classic endurance training adaptation in human muscle, this indicates the superiority of our workflow when it comes to metabolic maturation of the tissues.

We observed that certain manipulations selectively impacted endurance rather than maximal force generation or only affected endurance during certain conditions. For example, manipulating glycogen synthase, central to energy production during exercise [27,61,62] only affected endurance without affecting maximal force production. Furthermore, clenbuterol enhanced power output and improved endurance during glucose-rich conditions, while endurance during glucose starvation was impaired as expected based on findings from clinical trials and transgenic rodent studies of β_2_ agonism of skeletal muscle [47–49]. This resolution underscores the model’s capacity to tease apart mechanistic contributions to distinct aspects of muscle plasticity, such as force vs. endurance or energetic production vs. structural adaptation. Responses also diverged under metabolic stress, as evidenced by glucose-deprivation experiments where glycogen-dependent pathways became limiting. This sensitivity and resolution to detect interventions complement previous work by Madden et al., [18] that reported increased myotube size following 2 weeks of high clenbuterol (100nM) stimulation. Here, we detected acute effects on signaling already at 1nM and dose-dependent chronic effects with doses as low as 0.2nM being effective at increasing maximal force both in control conditions and following detraining. Furthermore, since β_2_ agonism via clenbuterol augmented contractile performance and preserved function during inactivity, we provide evidence of the utility of this platform to test pharmacological interventions relevant to muscle atrophy and anabolic resistance during physiologically relevant conditions such as inactivity-associated detraining rather than during atrophy induced by supraphysiological concentrations of e.g. Activins or glucocorticoids previously used by us [63] and others [24].

Earlier reports on the effect of exercise training-mimicking stimulation regimens on acute fatigue development have been inconsistent [19][24]. The interaction of the fatigue protocols to physiological interventions like hypoxia and glucose starvation have not been assessed, making interpretation of the data difficult. Here, we confirmed improved endurance following repeated stimulation in accordance with Pallotta et al. [24] and report robust and expected response to both hypoxia and glucose starvation.

Standard 2D human myotube cultures are limited as experimental models to study insulin-regulated glucose metabolism because they depend heavily on GLUT1 and GLUT3, express low levels of GLUT4, and show weak insulin-stimulated GLUT4 translocation to the plasma membrane [64]. In our system, we observed robustly increased GLUT4 expression and a GLUT4 spatial pattern resembling adult human muscle [52–54,65]. These features are an improvement over prior human 2D cultures as well as in tissue-engineered myobundles [66]. By promoting physiologically relevant GLUT4-dominant expression and regulation, this model likely provides a more robust representation of human muscle insulin action and a valuable tool for both insulin and GLUT4 biology research and drug discovery.

Finally, this work reveals causality for GYS1 in sustaining human muscle endurance. Silencing GYS1 did not affect baseline maximal tetanic force production or fatigue resistance under standard culture conditions but severely compromised endurance capacity when glucose availability was restricted. These findings highlight that glycogen metabolism is dispensable for acute maximal force production but becomes essential under contraction-induced fatigue development in conditions where the other major glycolytic substrate glucose is unavailable. This supports the notion that GYS1 confers metabolic resilience by maintaining internal energy reserves. This has implications for understanding human diseases characterized by glycogen dysregulation, such as McArdle disease, and may inform on therapeutic strategies targeting muscle metabolism in aging, insulin resistance, and inactivity.

In conclusion, our study establishes a functionally mature, plastic, and genetically tractable human muscle workflow that faithfully models both short- and long-term adaptations to contractile activity and metabolic stress. The ability to integrate physiological stimuli, pharmacological agents, and targeted gene manipulation opens new avenues for studying human muscle biology and advancing therapeutic discovery in metabolic and neuromuscular disease contexts.

## 4. MATERIALS AND METHODS

### Bioengineered muscle tissue formation

Three-dimensional bioengineered muscle tissues were produced based on primary myoblasts from a healthy donor (Cook Myosite) and from leveraging a previously described protocol [15]. 300,000 cells were resuspended in 43µl Hams F10 medium (Gibco), supplemented with 12µl Matrigel, 4µl 50mg/mL fibrinogen (Sigma-Aldrich), and 1.2µl 100U/µl thrombin (Sigma-Aldrich) for a total seeding volume of 60µl, 5 million cells per ml. The cell suspension was applied to a Mantarray casting well with mini-sized wells and a two-post array (Curibio) and incubated at 37°C, 5% CO2 for 80 min for hydrogel formation. Next, 1 mL of F10 medium was added to the wells and after an additional 10-minute incubation the posts were transferred to Myotonic medium (Cook Myosite) supplemented with 5g/mL aminocaproic acid (Sigma-Aldrich). 24 hours after casting the bioengineered muscle tissues were transferred to Primary Skeletal Muscle Differentiation medium (#SKM-MED-PRI-D, Curibio). Medium was refreshed every 2-3 days throughout experimentation. After 7 days, the bioengineered muscle tissues were transferred to Primary Skeletal Muscle Maintenance Medium (SKM-MED-PRI-M Curibio) and contractile assessment was initiated. For experiments assessing glucose uptake and glycogen synthesis the bioengineered muscle tissues were cultured in metabolic differentiation medium (#META-PRI-250-D, Curibio) from day 0-10 and then transferred to metabolic maintenance medium (#META-PRI-250-M, Curibio) instead of maintenance medium. Tissues were cultured for up to a total of 6 weeks as indicated in the figures.

### Contractile analyses

#### Tissue maturation monitoring and maximal force assessment

Ahead of each medium change force production in the bioengineered muscle tissues was assessed. Assessment was performed using a purpose-built cell culture plate lid supporting graphite electrodes compatible with the Mantarray hardware (Curibio). Force was evaluated by applying biphasic 75 mA pulses of 10ms for 2 seconds at 1, 2, 3, 5, 10, 20, 30, and 40Hz, with 8 seconds between each frequency. This was repeated twice. Twitch force was quantified as the mean of the two twitches in the last stimuli and maximal tetanic force was measured as the peak force during the last 40Hz stimulation. This assessment was used to monitor the maturation of tissues as well as read out the maximal force.

#### Fatiguing contraction bout and recovery

To assess endurance and recovery capacity of the tissues two different contraction protocols were applied as noted in the figure legends. Protocol 1 consisted of first 5 minutes of tetanic contractions (50Hz, 5ms, 100mAmp, Biphasic, 0.5 sec) every 2 sec and then 4 minutes of tetanic contractions every 1.5 sec. Protocol 2 consisted of contractions every 2 sec throughout. Following the last contraction, tissues were allowed to recover for 20 sec. before three additional tetanic stimulations were applied. A trajectory of the peak from each individual tetanus was calculated and from this the area under the curve was calculated as a measure of endurance. The ratio between the average force production from the first 3 tetani and the average force production from the 3 tetani during tissue recovery was calculated as a measure of recovery ability.

#### Exercise-mimicking electrical stimulations of tissues

Exercise training mimicking contractions were evoked by stimulating the tissues (50Hz, 5ms, 100mAmp, Biphasic, 5 sec every 50 seconds), first, 1×30 min, then 1×60 min and then continuing 1×90 min stimulation throughout the intervention. Stimulations were performed Monday, Wednesday, Friday.

### Sample lysing, western blotting and qPCR

Tissues were harvested by removing them from their posts, blotting them on filter paper and snap freezing them in liquid nitrogen. Next tissues were homogenized by placing the frozen tissue in a 2ml Eppendorf tube with a steel bead and 300µl RLT buffer containing 40µM DTT (for RNA extraction) or homogenization-buffer (10% glycerol, 20 mM Na_4_P_2_O_7_, 150 mM NaCl, 50 mM HEPES, 1% Tergitol 15-S-40, 20 mM beta-glycerophosphate, 10 mM NaF, 2 mM PMSF, 1 mM EDTA, 1 mM EGTA, 3 mM benzamidine, 10 µg/mL leupeptin, 10 µg/mL aprotinin, 2 mM Na_3_VO_4_, pH=7.5) (for western blotting) and shaken in a bead mill tissue lyser at 30Hz for 60sec. Subsequently, samples were spun at 10.000rpm, 4 C for 20 min and super natant was transferred to new tubes and frozen until further analyses.

For qPCR and RNAseq analyses, extraction was performed with RNeasy Fibrous Tissue Mini Kit (Qiagen 74704) according to manufacturer’s protocol. For cDNA synthesis iScript cDNA Synthesis kit (Bio Rad 1708891) was used. The following primers were used: TaqMan Gene Expression Assays (ThermoFisher Cat# 4331182), with the following ID numbers: GYS1: ID Hs00157863_m1, Ppargc1a: ID Hs00173304_m1, Nr4a3: ID Hs00545009_g1, Nr4a1: ID Hs00374226_m1 and Pgk1: ID Hs00943178_g1. For RNAseq the library prep and sequencing were performed by Lexogen.

For western blotting, lysate protein concentration was determined using the bicinchoninic acid method, and immunoblotting was performed as previously described [67]. The following antibodies were used: SERCA2 (SCBT sc-53010), Hexokinase II (SCBT sc-130358), Glycogen synthase (was a kind gift from Oluf Pedersen, University of Copenhagen, DK), Phosphofructokinase (SCBT sc-166722), Mito cocktail (NDUFB8, SDHB, UQCRC2, COX-II, ATP5A) (Abcam ab110411), Citrate Synthase (Abcam ab96600), AMPK α_1_ (a kind gift from Olga Göransson, Lund University, SE), α_2_ (MRC PPU Reagents and Services, University of Dundee, Scotland, UK, custom made), β_1_ (SCBT sc-100357), β_2_ (1.5, a kind gift from Dr. DG Hardie, University of Dundee, Scotland, UK), γ_1_ (Abcam ab32508), γ_3_ (Genscript, NJ, USA, custom made), Myogenin (Abcam ab1835), MyoD1 (Thermo Scientific, Waltham, MA, PA5-23078), pHSL Ser660 (CST #4126), pCREB Ser133 (CST #9198), pRPS6 Ser 236/236 (CST #2211), pRPS6 Ser240/244 (CST #2215), GLUT4 (Thermo Scientific, Waltham, MA, PA1-1065), PDH-E1α (SC-377092). Myosin Heavy Chain (MHC) composition was determined with SDS-PAGE separation in Stain-free gels as described previously [68].

### RNAseq data analyses

3’ library (QuantSeq) preparation and sequencing (SE100), sequence alignment, count quantification and data QC were performed by Lexogen. Briefly, raw sequencing reads were evaluated with FastQC, adapters and low-quality bases were trimmed with Cutadapt, and MultiQC was used to aggregate QC across samples for both raw and trimmed reads. The trimmed reads were then aligned with STAR [64] (Spliced Transcripts Alignment to a Reference) to the GRCh38 (release 94) reference genome augmented with ERCC (External RNA Controls Consortium) and SIRV (Spike-In RNA Variant) sequences. Post-alignment QC was provided by STAR’s mapping summaries, and RSeQC read-distribution analyses profiled tags across CDS, UTRs, introns, and intergenic regions. Reproducibility was evaluated via inter-replicate correlation using orthogonal regression within each condition. Based on the post-alignment QC, all 22 samples can reasonably be considered to have passed. Differential gene expression analysis between various time points and conditions were performed using DEseq2 [62]. *P* values were corrected for multiple testing using Benjamini and Hochberg method. Adjusted *P* value < 0.05 was used as a cutoff for differentially expressed genes (DEGs). Over-representation analysis (ORA) was performed for DEGs using clusterProfiler [63] to find enriched GO terms. Time-course DEGs were clustered with Mfuzz [61]; the number of clusters was selected from an elbow plot of the minimum centroid distance across 2-10 clusters. ORA was performed for genes in each cluster to examine biological relevance. Adjusted *P* value < 0.05 was used as a cutoff for significant enrichment.

### Imaging analyses

Muscle tissue were fixed with 4% paraformaldehyde (PFA) diluted in PBS at room temperature for 15 minutes, then washed three times for 5 minutes in PBS. Bundles were permeabilized with 0.1% Triton X-100 for 20 minutes, followed by an additional series of PBS washes. Samples were transferred into blocking buffer containing 1% BSA for 30 minutes to minimize nonspecific antibody binding. After blocking, bundles were incubated with primary antibodies diluted in blocking solution for 4 hours at 4 °C with gentle agitation. Following incubation, samples were washed several times in PBS to remove unbound antibodies. Bundles were then incubated with the appropriate fluorophore-conjugated secondary antibodies for 30 minutes in the dark. Nuclear staining with DAPI and phalloidin dyes was performed during secondary antibody incubation. After final PBS washes, tissues were mounted using [mounting medium, catalog number] on glass-bottom dishes or imaging chambers suitable for confocal microscopy. Tissues were imaged on a LSM 980 equipped with Airyscan detection, using 5× and 63× objectives. Images were processed using identical parameters across samples.

### Glucose uptake assessment

Skeletal Muscle tissue bundles were incubated in starvation medium (1× DMEM (Gibco #A1443001) supplemented with 1% penicillin/streptomycin and 110 mg/L sodium pyruvate (Gibco #11360039) for 2 h at 37 °C, 5% CO₂. Then bundles were transferred to a new media plate ±insulin for 1h and then moved to a new plate containing 2DG (Sigma-Aldrich #D3179) for an additional 30-min incubation. Next bundles were briefly blotted on filter paper and snap-frozen in liquid nitrogen. Subsequently tissues were lysed in 50 µL lysis buffer (1.5% Triton X-100 in DPBS [Gibco #14190-094]) and incubated on a 96 °C heating block in 25 µL Stop Buffer (Promega kit, *#J1343*) with intermittent vortexing until they were visually dissolved. Then 25 µL Neutralization Buffer (Promega kit, *#J1343*) were mixed in at room temperature. Homogenates were centrifuged at 18,000g for 4 min at 4 °C. For detection, homogenate was mixed 1:1 with prepared (1h prior) 2DG6P detection reagent and incubated for 60 min at room temperature. After 1h equilibration luminescence was recorded.

### GYS1 gene silencing

RNA strands for siRNA duplexes were synthesized purified at Biosynthesis (Lewisville, TX). The GYS1 siRNA had a 36-mer and 22-mer duplex RNA structure, with a 36-nt sense strand composed of modified RNA and a C22 lipid conjugation that was annealed to a 22-nt modified RNA antisense strand After 21 days of differentiation, GYS1 siRNA was added directly to the medium for a final concentration of 100nM. A matching volume of PBS was added in the control group. 3 days later the medium was changed to fresh medium without siRNA.

### Statistical testing

Results are shown as mean ± standard error of mean (SEM) with individual values shown on bar graphs and mean ± SEM for curves. Statistical testing was performed using t-test, one-way or two-way (repeated measures when appropriate) ANOVA as applicable. Tukey’s multiple comparison test was performed for post hoc analyses. Statistical analyses were performed using GraphPad Prism 10 (GraphPad Software, La Jolla, CA, USA, RRID: 002798). The significance level for all tests was set at α < 0.05.

## 5. Contributions

CHO and JRK conceived the study and wrote the initial draft. CHO, MKH, CTAL, JB, PJ, PB, BFH, RMV, JH, ZW, CSW, JW, TEJ, AB, CP and JRK performed and/or conceptualized the experiments. CHO, CTAL, TW, ZL, ET, and JRK performed data analyses and generated figures. JW, TEJ, AB, CP and JRK secured resources. All authors edited and reviewed the manuscript.

## 6. Acknowledgement

We thank the team at Skeletal Muscle Biology, Novo Nordisk for fruitful discussions and feedback on this work. We also thank Julia Prats for the support with staining of the muscle bundles. We acknowledge the Core Facility for Integrated Microscopy, Faculty of Health and Medical Sciences, University of Copenhagen.

## 7. Funding

JRK were supported by a grant from the Danish Diabetes Academy supported by Novo Nordisk Fonden (17SA0031406). JFPW and JBB was supported by grants from Novo Nordisk Foundation (NNF 082659, NNF 0085866)

## 8. Competing interests

CHO, MKH, CTAL, TW, ZL, PJ, PB, BFH, ET, JH, ZW, CSW, AB, CP and JRK were employed at Novo Nordisk A/S while the experimental work took place. JFPW is shareholder in and is external consultant for Novo Nordisk Inc.

---

**Figure S1.**
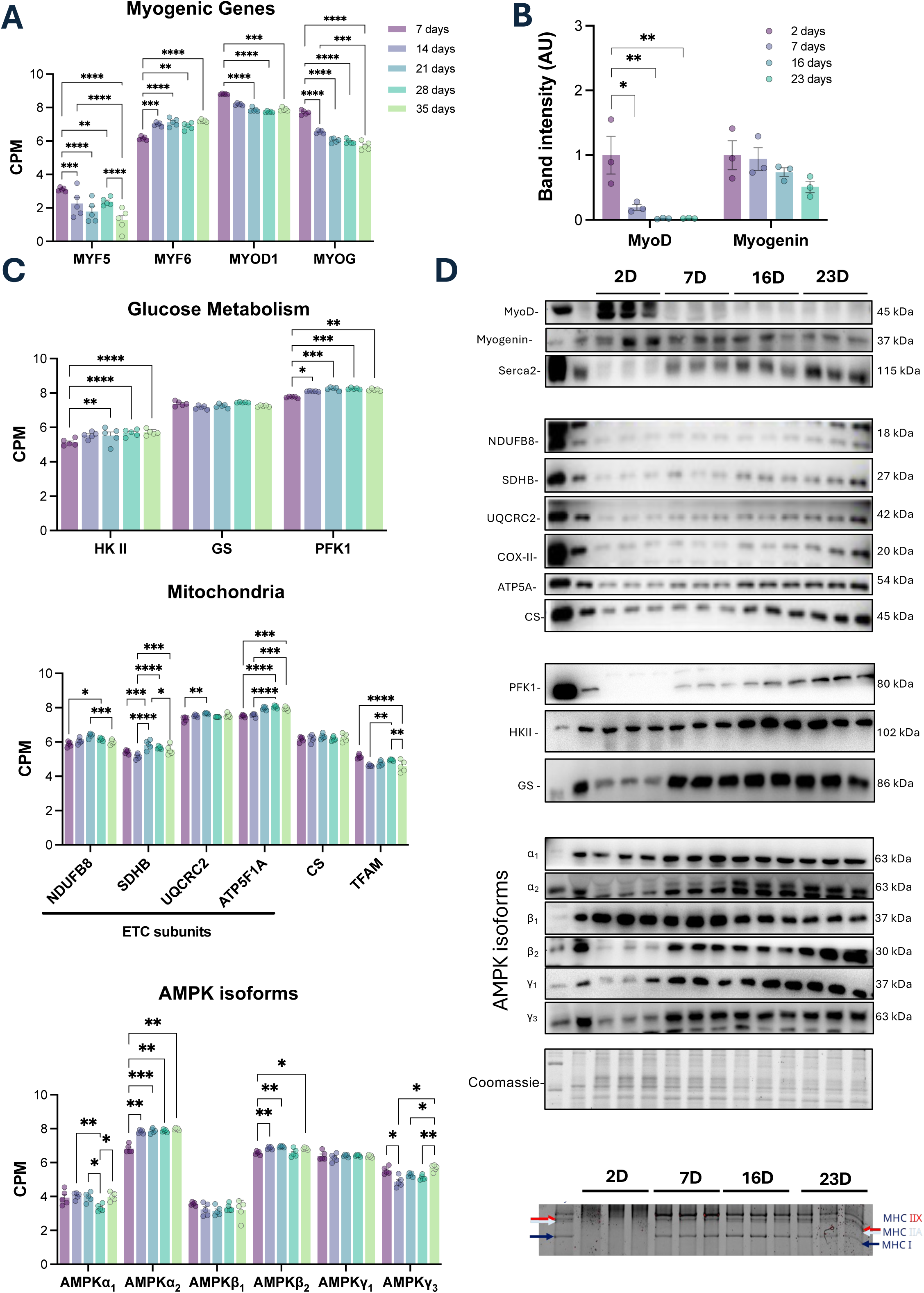
Transcriptional and protein validation of structural and metabolic maturation in bioengineered human muscle tissues. **(A)** RNA-seq expression (counts per million, CPM) of canonical myogenic regulators (MYF5, MYF6, MYOD1, MYOG) across the 3D tissue maturation time course (days 7, 14, 21, 28, and 35). **(B)** Quantification of MYOD and myogenin protein abundance across maturation (days 2, 7, 16, and 23) from immunoblots in D. **(C)** RNA-seq expression (CPM) of metabolic maturation markers grouped by pathway: glucose metabolism (GYS1, HK2, PFKM), mitochondria/oxidative phosphorylation (ATP5F1A, CS, NDUFB8, SDHB, TFAM, UQCRC2), and AMPK subunit isoforms (PRKAA1/2, PRKAB1/2, PRKAG1/3) across the 3D maturation time course (days 7–35). **(D)** Representative western blots across maturation (days 2, 7, 16, and 23) for myogenic regulators (MYOD, myogenin), contractile/transport proteins (SERCA2), mitochondrial/OXPHOS proteins (NDUFB8, SDHB, UQCRC2, COX-II, ATP5A, CS), glycolysis (PFK1, HKII), glycogen metabolism (GS), and AMPK subunit isoforms (α1, α2, β1, β2, γ1, γ3). Coomassie staining is shown as a loading control. Bottom panel shows myosin heavy chain isoforms (MHC I, MHC IIa, MHC IIx). Statistical comparisons were performed using one-way ANOVA with appropriate multiple-comparisons correction. \**p* < 0.05, \*\**p* < 0.01, \*\*\**p* < 0.001, \*\*\*\**p* < 0.0001.

**Figure S2.**
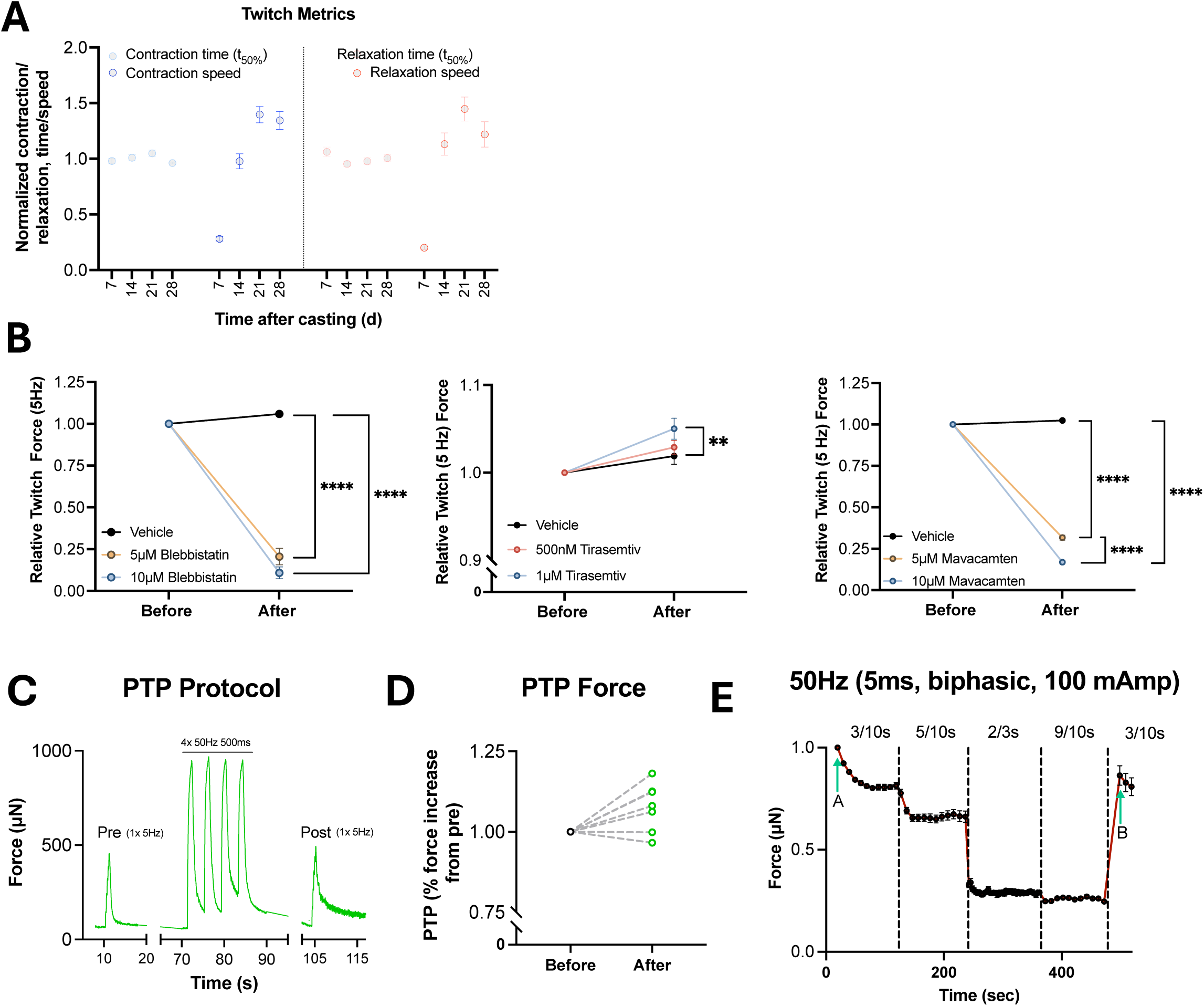
Contractile kinetics, pharmacological modulation, post-tetanic potentiation, and fatigue protocols in bioengineered human muscle tissues. **(A)** Normalized twitch kinetics across maturation, including contraction time, relaxation time, contraction speed, and relaxation speed. **(B)** Fiber-type and cross-bridge modulation of force. Relative twitch force (5 Hz) measured before and after acute treatment with the slow-twitch myosin inhibitor mavacamten (5 or 10 µM), the fast-skeletal troponin activator tirasemtiv (500 nM or 1 µM), or the pan-myosin inhibitor blebbistatin (5 or 10 µM). **(C)** Representative trace of the post-tetanic potentiation (PTP) protocol: a baseline twitch (pre), followed by four tetanic trains (50 Hz, 500 ms), and a post-tetanic twitch (post). **(D)** Quantification of PTP expressed as percent increase in twitch force after tetanic stimulation relative to pre-tetanus twitch force (paired measurements per tissue). **(E)** Representative force trace illustrating the stepwise fatigue protocol based on repeated 50 Hz tetanic stimulation (5 ms biphasic pulses, 100 mA) with escalating cycles. Statistical comparisons were performed using t-student, one-way or two-way ANOVA with appropriate multiple-comparisons correction. \*\**p* < 0.01, \*\*\**p* < 0.001, \*\*\*\**p* < 0.0001.

**Figure S3.**
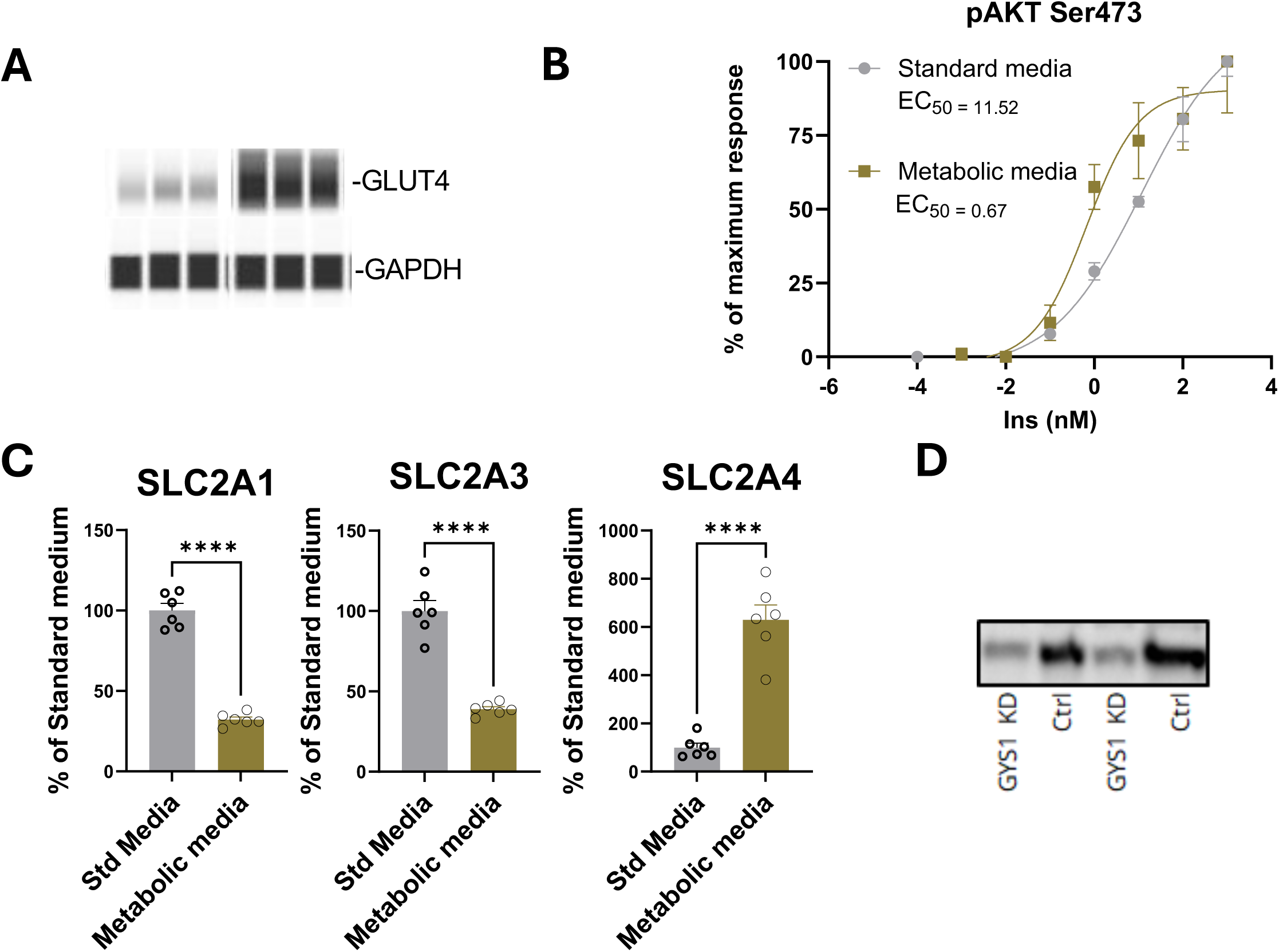
Medium optimization improves insulin signaling and glucose transporter expression, and representative validation of GYS1 knockdown. **(A)** Insulin dose–response of Akt phosphorylation at Ser473 (pAKT Ser473) in 7-day differentiated monolayer myotubes cultured in standard versus metabolic media. Data are expressed as % of maximal response; EC50 values are indicated. **(B)** Relative mRNA expression of glucose transporter genes SLC2A1 (GLUT1), SLC2A3 (GLUT3), and SLC2A4 (GLUT4) in myotubes cultured in standard versus metabolic media. **(C)** Representative western blot showing GYS1 protein levels in tissues subjected to GYS1 knockdown (GYS1 KD) versus control (Ctrl). Bars show mean ± SEM with individual values. Statistical comparisons were performed using t-student, ****P < 0.0001.

